# *Lactococcus lactis sb. cremoris* orchestrates signal events in the gut epithelium via TLR2 to promote tissue restitution

**DOI:** 10.1101/2021.12.02.471025

**Authors:** Crystal R. Naudin, Joshua A. Owens, Lauren C. Askew, Ramsha Nabihah Khan, Christopher D. Scharer, Jason D. Matthews, Liping Luo, Jiyoung Kim, April R. Reedy, Maria E. Barbian, Rheinallt M. Jones

## Abstract

The use of beneficial bacteria to promote gastrointestinal heath is widely practiced, however, the mechanisms whereby many of these microbes elicit their beneficial effects remain elusive. Previously, we conducted a screen for the discovery of novel beneficial microbes and identified the potent cytoprotective effects of a strain of *Lactococcus lactis* subsp. cremoris. Here, we show that dietary supplementation with *L. lactis* subsp. cremoris induced transcript enrichment of a set of genes within the colon whose functions are associated with host cell and microbe interactions. Specifically, *L. lactis* subsp. cremoris induced the expression of *tlr2*, which we show was required for *L. lactis* subsp. cremoris to elicit its beneficial effects on the intestine. *L. lactis* subsp. cremoris did not confer beneficial effects in mice deficient in TLR-2, or deficient in its adaptor protein Myd88 in chronic gut injury models. In addition to cytoprotection, culture supernatant from *L. lactis* subsp. cremoris accelerated epithelial migration in a cultured epithelial cell scratch wound assay; and effect that was abrogated by a TLR-2 antagonist. Furthermore, *L. lactis* subsp. cremoris accelerated epithelial tissue restitution following the infliction of a colonic wound biopsy in a TLR-2 and Myd88-dependent manner. Within colonic wounds, *L. lactis* subsp. cremoris induced the activation of signaling pathways that function in tissue restitution following injury, including the ERK signaling pathway, and of focal adhesion complex (FAC) proteins. Together, these data demonstrate that *L. lactis* subsp. cremoris signals via the TLR2/MyD88-axis to confer cytoprotection and accelarated tissue restituion in the gut epithelium. These data point to evolving adaptations where beneficial gut microbes moduate innate immune signaling to excert positive influnces on host physiology.

## Introduction

Supplementation of the diet with live bacteria, termed probiotic therapy, is a continuously expanding market, with recent reports recording that up to 4 million adults consume probiotics in the USA, and with up to 60% of healthcare providers prescribing probiotics to patients [1, 2]. Consumption of probiotics has been reported to improve resistance against pathogens [3], prevent cardio-metabolic disease [4, 5], help mental and behavioral wellbeing [6], ease gastrointestinal (GI) distress [7, 8], and priming of the immune system [9, 10]. In spite of these reports, evidence of probiotic efficacy remains sparse, especially in humans [11, 12], and the efficacies of some probiotics for existing conditions is highly debated [13]. Recent reports have examined the extent to which probiotics colonize the human GI tract, and in particular the mucosa-associated surfaces [14]. In addition, the modulatory effects of probiotics on community structure of the resident microbiome in people have not been thoroughly examined, with a recent review reporting no effect of probiotics on fecal microbiome composition [15], and other studies observing modifications in the fecal microbial community structure in patients that ingested probiotics [16–18]. Due to the shortage of conclusive empirical evidence, further investigations focused on the effects of probiotics on the host are required for clinicians and consumers to be able to make full and educated choices regarding therapies.

Inflammatory bowel disease (IBD) is caused by defects that interact across a genetic, immunologic and environmental axis, creating symptoms that are often difficult to treat [19]. Disease of the intestine including Crohn’s disease (CD) and Ulcerative colitis (UC), have at their source, uncontrolled gut mucosal inflammation [20, 21]. The intestine is lined by epithelial cells that form a strong yet selectively permeable barrier to protect the underlying lamina propria from luminal contents. However, sustained inflammation experienced during IBD often leads to increased gut permeability and breaches in the epithelial barrier that can often exacerbate damaged mucosa [21, 22]. A delay in promoting intestinal barrier restitution can cause further tissue damage that increase wounding and leads to chronic disease [20, 21, 23]. The chronic inflammation experienced by IBD patients often disrupts the normal healing process and creates defects in wound repair [24]. A decrease in inflammation is a primary therapeutic goal for most IBD patients, and generally a first line treatment. Since the intestinal epithelium is paramount to wound healing, augmenting the signaling pathways that drive reorganization of the epithelium and promote the growth and repair necessary to rebuild the intestinal mucosa is critical.

The microbial composition in IBD patients becomes aberrant with normal microflora such as the genera *Bifidobacterium* and *Lactobacillus* decreasing, while pathogenic or potential harmful bacteria are increased [25, 26]. Critically, dietary supplementation with beneficial microbes may balance the indigenous microbiome in IBD patients, and have a therapeutic effect on IBD [27, 28]. Indeed, beneficial microbes, that induce an advantage to health [29, 30] can prompt a change in the diversity and metabolic activity of the gut microbiota [31], modulate innate and adaptive immune responses [32, 33], or significantly improve epithelial barrier function [34, 35]. Despite this knowledge, the lack of empirical data to substantiate claims made by marketed microbial products poses a considerable challenge to consumers and caregivers when selecting a suitable microbe as a potential therapeutic intervention to treat a given ailment, including IBD. Thus, there is a pressing need to expand primary scientific literature that experimentally substantiates the proposed uses of beneficial microbes.

Our group has reported on the conserved molecular mechanisms of host cell and beneficial microbe interactions in the intestine of *Drosophila* and mice [36–38]. Using assays we developed to show the mechanism of the beneficial effects of microbes in the Drosophila gut, we initiated a screen to discover new and potentially more efficacious beneficial bacteria. To this end, we successfully identified a previously uncharacterized beneficial bacterium, namely *L. lactis* subsp. *cremoris* ATCC19257 that elicited highly efficacious protective influences on the *Drosophila* and mouse intestine [39]. The objective of this study is to identify the key molecular components and functional elements that mediate *L. lactis* subsp. cremoris capacity to promote the host’s ability to restrain intestinal inflammation and accelerate restitution of intestinal injury and wound repair. These studies will contribute to inform potential ‘Early Clinical Trial with Live Bio-therapeutic Products’, using *L. lactis* ATCC19257 to treat intestinal diseases. To identify the molecular mechanisms whereby *L. lactis* subsp. cremoris elicits its beneficial effects on the host, we characterized colonic gene expression induced by *L. lactis* subsp. cremoris, the host sentinel elements that recognize *L. lactis* subsp. cremoris, and the downstream host signaling pathways that are activated to elicit beneficial influences.

Herein, we show that *L. lactis* subsp. cremoris induced the expression of a set of genes in the colon that are associated with host tissue homeostasis and response to injury, including the JAK-STAT signaling pathway. Kegg pathway analysis also identified TLR2 and Myd88 as potential mediators of host cell and *L. lactis* subsp. cremoris signaling. Indeed, we show that *L. lactis* subsp. cremoris also elicited cytoprotection against DSS experimentally induced chronic colitis injury in mice in a Myd88 and TLR2-dependent manner. Furthermore, the culture supernatant of *L. lactis* subsp. cremoris induced cell proliferation in both cultured human epithelia cells, and in ex vivo organoids and TLR2-dependent manner. The capacity to accelerate wound restitution is a valuable property of putative beneficial bacteria. We show that the filtered supernatant from cultures of *L. lactis* subsp. cremoris was sufficient to accelerate restitution of wounds inflicted on cultured cells, and also show that supplementation of *L. lactis* subsp. cremoris accelerated the restitution of or on biopsy wound inflicted on colonic tissue. In both cultured cells and colonic tissue, *L. lactis* subsp. cremoris activated the expression of Focal adhesion complex (FAC) proteins which function in cell movement on the leading edge of healing wounds in a TLR2 - dependent manner. These data show that *L. lactis* subsp. cremoris signals via the TLR2/MyD88-axis to confer cytoprotection and accelarated tissue restituion in the gut epithelium. These data show that beneficial *L. lactis* subsp. cremoris can moduate innate immune signaling to confer beneficial influnces on host physiology.

## Results

### *L. lactis* subsp. cremoris dietary supplementation induces transcript enrichment within the colon

To assess the extent to which *L. lactis* subsp. cremoris activates gene expression in the colon, germ-free C57BL/6 mice were fed 1×10^8^ CFU *L. lactis* subsp. cremoris by oral gavage. 4 hours after ingestion, the mice were sacrificed, colons were harvested, and transcript enrichment measured by RNAseq analysis. Principle component analysis (PCA) analysis showed that the transcriptome of mice fed *L. lactis* subsp. cremoris clustered to a tight and separate group from control fed vehicle mice (**Figure 1A**). Representation of the transcriptome of these mice by volcano plot analysis identified the most statistical significant and highest fold change genes activated in the colon by *L. lactis* subsp. cremoris (**Figure 1B**). Visualizing the expression of genes across the samples by heat map analysis identified the top differentially expressed genes in the colon activated by *L. lactis* subsp. cremoris, including the upregulation of *tlr2* transcript levels (**Figure 1C**). In addition, KEGG pathways based on all differentially expressed genes revealed significantly enriched activation of cell signaling pathways that function in host tissue homeostasis, including the JAK-STAT pathway (**Figure 1D and 1E**). These data are consistent with our previous report where we showed JAK-STAT pathways signals can be initiated by probiotic lactobacilli in the gut lumen to sub-epithelial compartments [40]. To establish how these pathways may be increased by *L. lactis* subsp. cremoris, we performed string pathway analysis using the leading edge genes identified in figure 1E and identified proteins that function in host cell and microbe interactions, including Myd88 and TLR2 (**Figure 1F**). These data show that *L. lactis* subsp. cremoris activates a specific genetic signature through host cell microbial sentinel receptors, which may mediate its beneficial influences on host health and disease.

**Figure 1:**
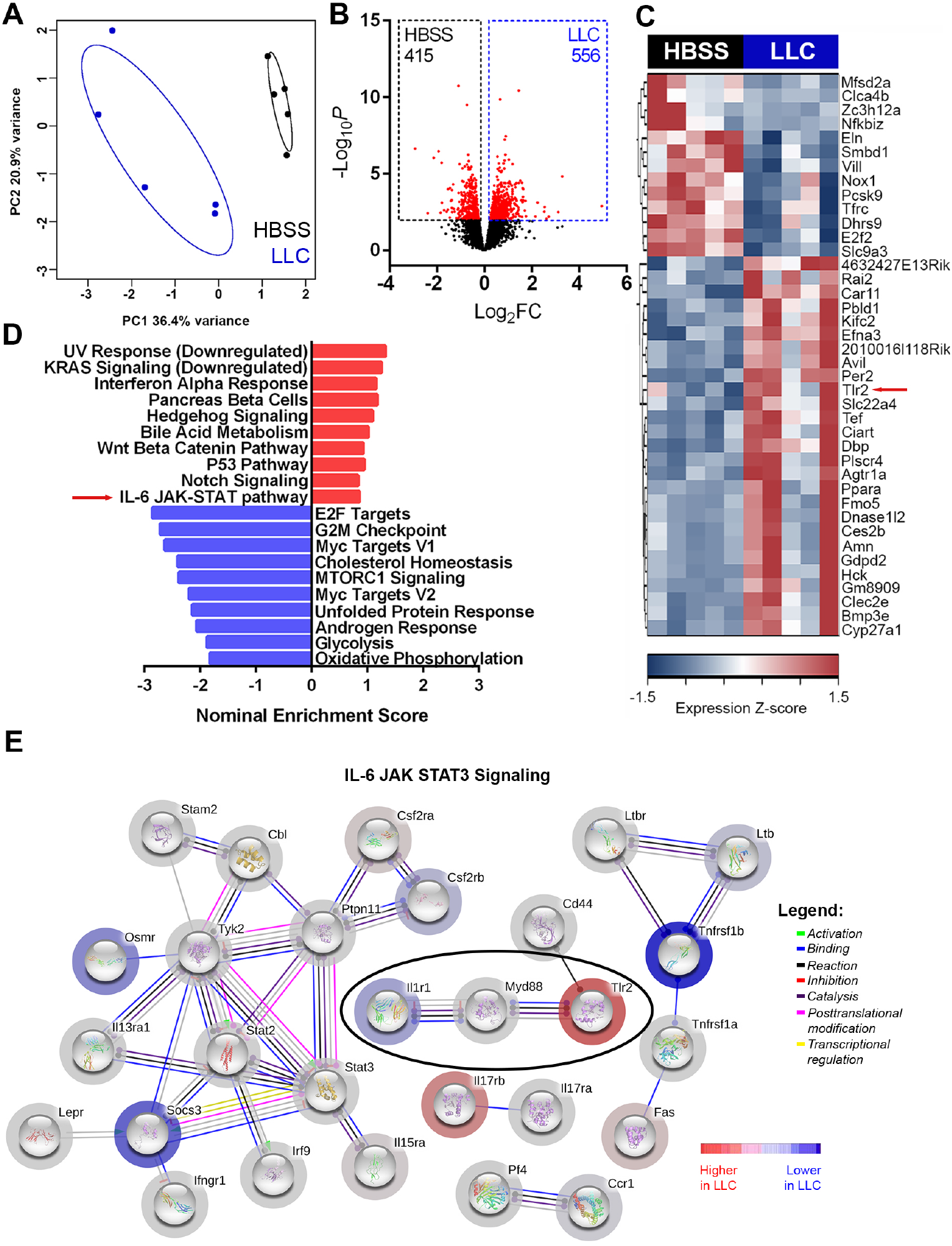
*L. lactis* subsp. cremoris treatment upregulates modulates IL-6 JAK-STAT gene expression in intestinal epithelium. **(A)** Principle component analysis (PCA) of mouse proximal colon transcriptome after LLC treatment. n = 5 mice per group. Permutational multivariate analysis of variance (PERMANOVA). **(B)** Volcano plot of differential expression of genes in LLC versus vehicle treated in (A) Red dots, differentially expressed genes (DESeq2 Wald test, false discovery rate (FDR) < 0.1). **(C)** Heat map showing the top 40 most significantly downregulated or upregulated genes (low-expression genes with mean normalized counts in the bottom 20th percentile were excluded). **(D)** Significantly enriched KEGG pathways based on all differentially expressed genes. **(E)** STRING network visualization of the genes in IL-6 JAK-STAT pathway. Edges represent protein–protein associations. Disconnected nodes were excluded.

### *L. lactis* subsp. cremoris dietary supplementation protects against gut injury induced by Dextran Sodium Sulfate (DSS) in a Myd88-dependent manner

A key mediator of host cell and microbe interaction is the pattern recognition receptor (PRR) adaptor protein Myd88. Because string pathway analysis pointed to a potential role of innate singling, we investigated if Myd88 is required for *L. lactis* subsp. cremoris to elicit its beneficial influences on host health and disease. We supplemented the diet of C57BL/6 wild-type and Myd88-/- mice daily for 2 weeks with 1×10^8^ CFU *L. lactis* subsp. cremoris or vehicle control by oral gavage. Mice were then subjected to multiple rounds of 3% DSS in their disease activity recorded. We first show that C57BL/6 mice that had been supplemented with *L. lactis* subsp. cremoris exhibited significantly less disease activity index compared to C57BL/6 that were treated with vehicle control (**Figure 2A**). *L. lactis* subsp. cremoris supplementation significantly lowered diseases parameters in wild type mice, but not in Myd88-/- mice up to day 13 following inclusion of DSS in the drinking water (**Figure 2B**). The protective influences of *L. lactis* subsp. cremoris were most pronounces at day 5 following inclusion of DSS in the drinking water (**Figure 2C**). We repeated the experiment as described in figure 2A but sacrificed the experimental groups at day 5 following after inclusion of DSS in the drinking water. *L. lactis* subsp. cremoris supplementation significantly protected against DSS-induced colon reduced length in wild type mice, but not in Myd88-/- mice (**Figure 2D and E**). Before mice were sacrificed, they were subjected to colonoscopy analysis. *L. lactis* subsp. cremoris supplementation markedly protected wild type mice, but not Myd88-/- mice from DSS-induced tissue injury, as detected by visibly less tissue injury in the colonoscope view, and by histological analysis (**Figure 2D and E**). These data indicate that *L. lactis* subsp. cremoris supplementation protects against acute gut injury induced by acute Dextran Sodium Sulfate (DSS) in a Myd88-dependent manner.

**Figure 2.**
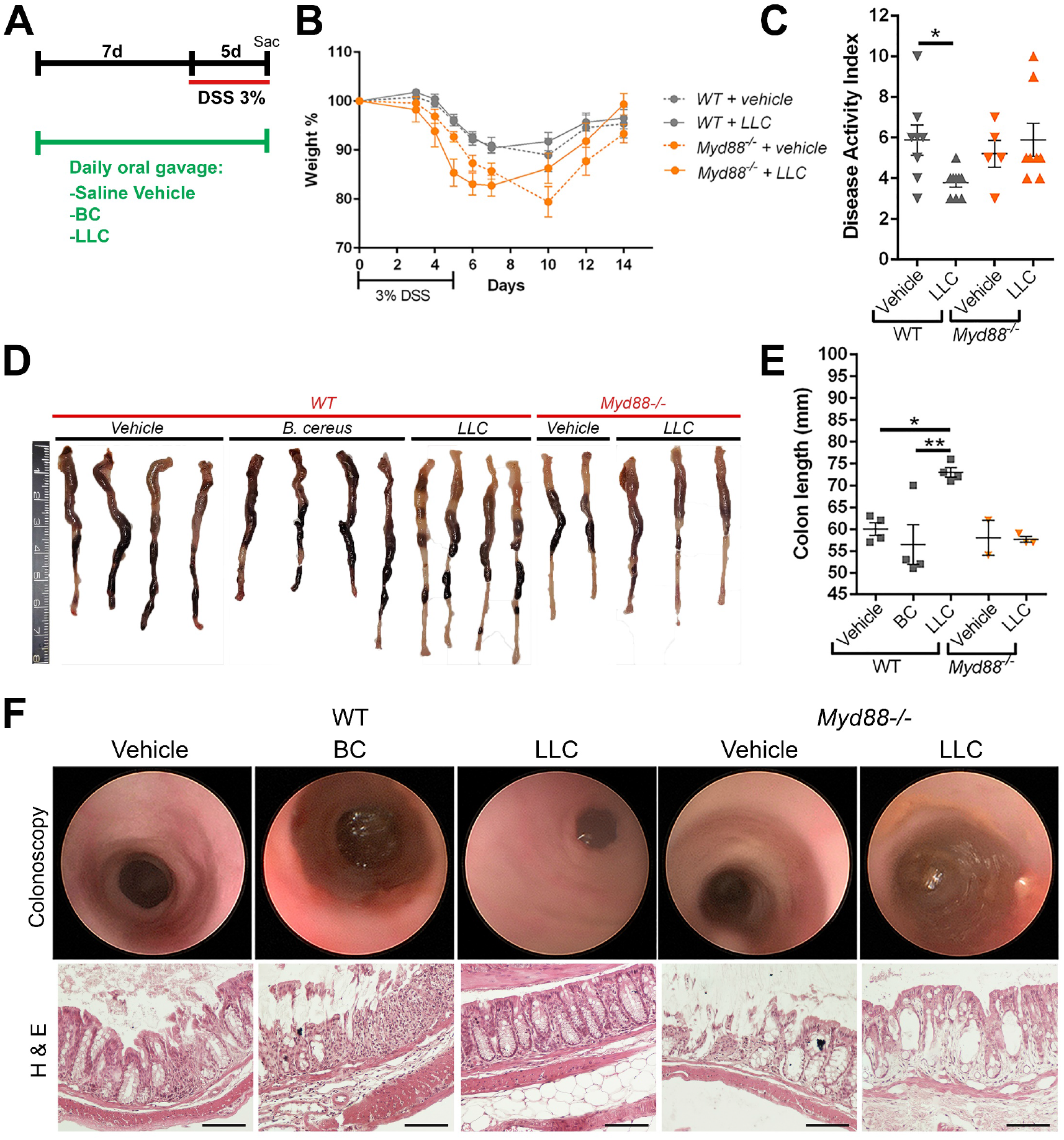
*L. lactis* subsp. cremoris treatment protects the murine intestine from acute Dextran Sodium Sulfate (DSS)-induced colitis in a Myd88 dependent manner. **(A**) Six-week-old male *Myd88-/-* and WT C57Bl/6 mice were administered either 1 × 10^9^ CFU total Lactococcus lactis subsp. cremoris ATCC 19257 (LLC), Bacillus cereus (BC) or HBSS (vehicle control) for 7 days. Daily probiotic gavage was continued daily as 3% DSS was included in the drinking water (day 0) for 5 days in the indicated groups. Weight was tracked in mice following commencement of the inclusion of 3% DSS in the drinking water. N = 8-10. **(B)** The body weights of all mice were monitored every 2-3 days. Weights and survival were tracked out to day 13 in a subset of mice with no significant difference between groups. **(C)** Disease activity index (DAI) of all mice in after 5 days of DSS treatment. **(D)** Colons of a subset of mice sacrificed after 5 days on DSS, N = 4, 4, 4, 2, 3 for groups: WT+ vehicle, WT+ BC, WT+ LLC, Myd88-/- + vehicle and Myd88-/- + LLC. **(E)** Colon length in millimeters, N = 4, 4, 4, 2, 3. **(F)** Representative colonoscopic images, and Swiss rolled mouse colons after 5 days of 3% DSS were stained with hematoxylin and eosin. Bars represent 500µM. Representative images are shown, N = 4, 4, 4, 2, 3. Data are represented as mean ± SEM, One way-ANOVA tests were used with multiple comparisons to determine the difference between groups *p < 0.05, **p < 0.01, ***p<0.001

### *L. lactis* subsp. cremoris supplementation protects against chronic DSS colitis and improves rate of healing in a Myd88 dependent manner

As an alternative to acute the DSS mouse injury model outlined in figure 2, we also assessed the extent to which *L. lactis* subsp. cremoris supplementation protects against the milder injury inducing chronic DSS colitis mouse model. Six-week-old C57BL/6 WT or *Myd88-/-* and mice were administered either 1 × 10^9^ CFU total *L. lactis* subsp. cremoris or HBSS (vehicle control) for 7 days before 2% DSS was included in the drinking water of experimental groups for repeated rounds over 28 days (**Figure 3A**). *L. lactis* subsp. cremoris supplementated mice had significantly lower disease activity in the colonic tissues in a Myd88-dependent manner over the entire study period (**Figure 3A-E**). At sacrifice at day 28, no differences were detected in spleen weight or colon weight between groups (**Figure 3F and G**). However, *L. lactis* subsp. cremoris supplementation significantly protected against DSS-induced reduced colon length, preserving colon length in wild type mice, but not in Myd88-/- mice at day 28 following the repeated DSS cycles. Furthermore, histological analysis of colonic tissue harvested at day 28 following the repeated DSS cycles confirmed that *L. lactis* subsp. cremoris supplementation mice significantly reduced colonic injury in a Myd88-dependent manner (**Figure 3I-M**). Together, these data show that *L. lactis* subsp. cremoris supplementation protects against chronic DSS colitis and improves healing rate following DSS injury of in a Myd88 dependent manner.

**Figure 3.**
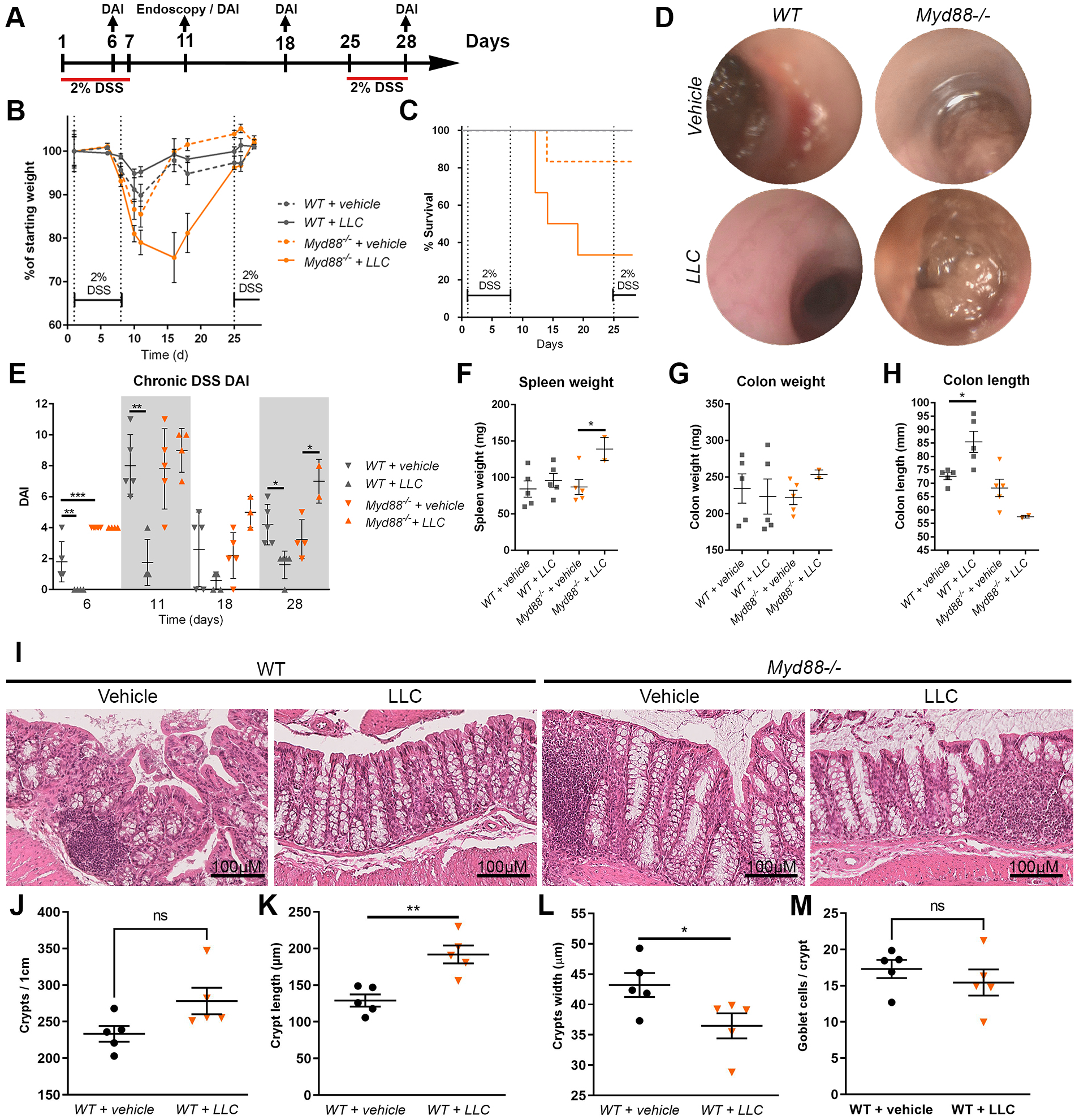
*L. lactis* subsp. cremoris treatment is beneficial against chronic DSS colitis and improves rate of healing in a Myd88 dependent manner. **(A)** Six-week-old male *Myd88-/-* and WT C57Bl/6 mice were administered either 1 × 10^9^ CFU total Lactococcus lactis subsp. cremoris ATCC 19257 (LLC) or HBSS (vehicle control) for 7 days. 2% DSS was included in the drinking water for 5 days in the indicated groups. Once all groups of mice had recovered to 95% of their original starting weights, 2% DSS was re-introduced into the drinking water for another 3 days. Weight was tracked in mice and probiotic or vehicle treatments were administered daily throughout. Data are represented as mean ± SEM. N=6. **(B)** The body weights of all mice were monitored every 2-3 days. **(C)** % survival. **(D)** Representative colonoscopic images at day 11, the mouse with median DAI on the day of colonoscopy in each group is shown. **(E)** Disease activity index (DAI) of mice at day 6, 11, 18 and 28. After 3 days on 2% DSS in the second round of DSS-induced colitis, surviving mice were sacrificed and **(F)** Spleen weight, **(G)** Colon weight and **(H)** Colon length were taken, N=6, 6, 5, 2. **(I)** Swiss rolled mouse colons were stained with hematoxylin and eosin (H&E, x200). Bars represent 100µM. A comparison of colon morphology was made between WT vehicle treated and WT LLC treated groups with regards to **(J)** crypts per cm, **(K)** crypt length **(L)**, crypt width and **(M)** number of goblet cells per crypt. Approximately 200 were measured per animal. N=5, 5. Data are represented as mean ± SEM. Data are represented as mean ± SEM, One way-ANOVA tests were used with multiple comparisons to determine the difference between groups *p < 0.05, **p < 0.01, ***p<0.001

### *L. lactis* subsp. cremoris promotes healing of biopsy wounds inflicted on the colon of in a Myd88-dependent manner

To confirm our observation of accelerated recovery following DSS-induced injury and faster wound healing in DSS injured mice, we inflicted repeated biopsies on the colon of individual mice and quantitated healing rates by colonoscopy. 6-week-old C57BL/6 WT or Myd88-/- mice were supplemented orally with either 1×10^8^ CFU total *L. lactis* subsp. cremoris, or with the non probiotic *B. cereus*, or vehicle control for 7 days. Mice were then inflicted with 4 wounds at distances of 1, 2, 3 and 4cm into the colon using biopsy forceps. Measurement of the rate of healing of biopsy was assessed by colonoscopy at 48 hours and 96 hours after initial wounding revealed that mice supplemented with *L. lactis* subsp. cremoris had significantly faster healing rates compared to mice supplemented with *B. cereus* or vehicle control (**Figure 4A-C)**.

**Figure 4.**
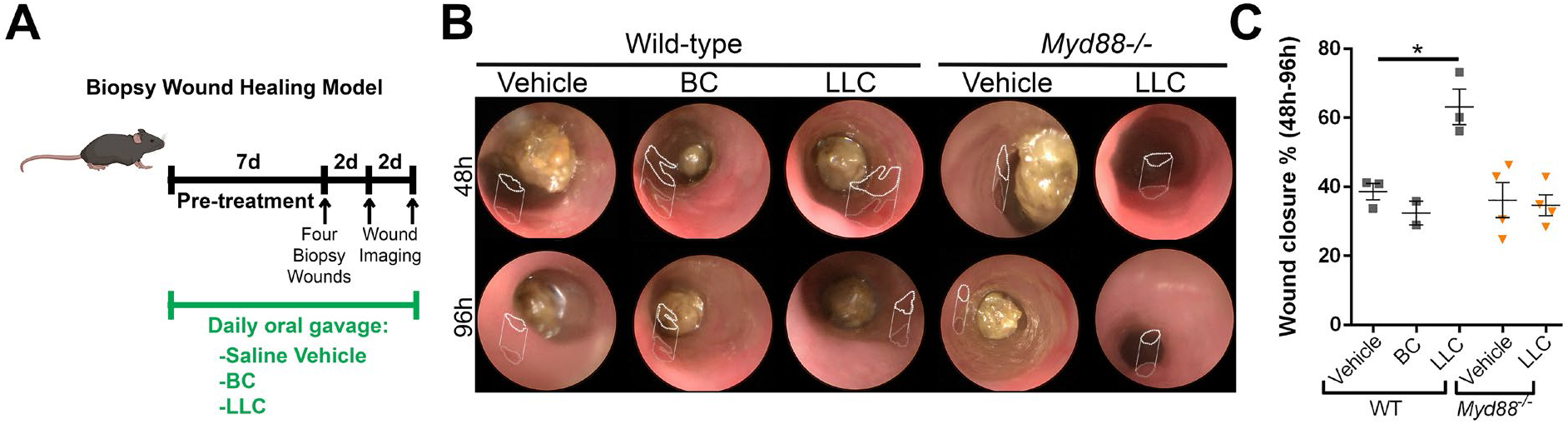
*L. lactis* subsp. cremoris treatment promotes wound healing in the colon in a Myd88 dependent manner. **(A)** Six-week-old male *Myd88-/-* and WT C57Bl/6 mice were administered either 1 × 10^9^ CFU total Lactococcus lactis subsp. cremoris ATCC 19257 (LLC), Bacillus cereus (BC) or HBSS (vehicle control) for 7 days. Mice were given four wounds at distances of 1, 2, 3 and 4cm into the colon using biopsy forceps. These wounds were tracked for healing using colonoscopy at 48 and 96h after initial wounding. N = 3-5. **(B)** Representative colonoscopy images. Wounds edges are indicated by dotted lines. **(C)** Percentage wound closure, each data point represents the average of four wounds in one mouse, data are represented as mean ± SEM, N = 2-5, One way-ANOVA tests were used with multiple comparisons to determine the difference between groups *p < 0.05, **p < 0.01, ***p<0.001.

Importantly, no differences in the healing rates of wounds were detected in Myd88-/- mice supplemented with *L. lactis* subsp. cremoris (**Figure 4A-C)**, showing that *L. lactis* subsp. cremoris supplementation promoted healing of biopsy wounds inflicted on the colon in a Myd88- dependent manner.

### *L. lactis* subsp. cremoris culture supernatant promotes cytoprotection and induces cell proliferation in cultured epithelial cells and in ex vivo cultured organoids in a TLR2-dependent manner

Dietary supplementation with *L. lactis* subsp. cremoris elicited beneficial influences on host colonic tissue. The colonic mucosa is separated from the luminal bacteria in the fecal stream by a thick mucus layer. Therefore, the beneficial bacteria do not come in direct physical contact with colonocytes, and therefore must exert their effects by the release of factors that penetrate the mucus layer. Upstream of Myd88 are the toll-like receptors (TLRs). Indeed, transcriptome analysis by RNAseq of colonic tissue of mice fed *L. lactis* subsp. cremoris detected an increase in expression of *tlr2* (**Figure 1C**). We confirmed this observation by quantitative (q)RT-PCR and show significantly enriched *tlr2* transcript levels in cultured T84 cells overlayed with media containing a 1 in 100 dilution of a filter sterilized culture supernatant collected from centrifuged cultures of *L. lactis* subsp. cremoris compared to media control (**Figure 5A**). To assess the extent to which *L. lactis* subsp. cremoris culture supernatant promotes wound healing, we inflicted a scratch wound on cultured T84 human epithelial cells, and then overlaid the cells with the filter sterilized culture supernatant of *L. lactis* subsp. cremoris or *B. cereus* or vehicle control at a 1:100 dilution factor for up to 72 hours and monitored healing rates. Culture supernatant of *L. lactis* subsp. cremoris significantly accelerated wound closure compared to *B. cereus* or vehicle control - a response that was abrogated by the presence of a TLR2 antagonist in the culture media (**Figure 5B-E)**. Cell movement is a critical aspect of wound healing that involves the migration of cells. Cell signaling networks including the ERK signaling pathway, and the composite of proteins that comprise the focal adhesion complexes (FAC) are mediators some of cell migration. We detected markedly increased levels of phosphorylated ERK in cultured cells overlaid with filter sterilized culture supernatant of *L. lactis* subsp. cremoris, compared to filtered media of *B. cereus* or vehicle control, or of filtered media from a culture of Lactobacillus rhamnosus GG that has been shown to activate ERK pathway signaling[41, 42] (**Figure 5F and 5G)**. We also detected markedly increased levels of phosphorylated focal adhesion kinase (p-FAK), a marker for FAC activity, in cultured cells overlaid with culture supernatant of *L. lactis* subsp. cremoris - a response that was abrogated by the presence of a TLR2 antagonist in the culture media (**Figure 5H)**. Together, these data show that culture supernatants promote healing of scratch wounds inflicted on cultured epithelial cells and activate cell signalling pathways that function in cell migration in a TLR2-dependent manner.

**Figure 5.**
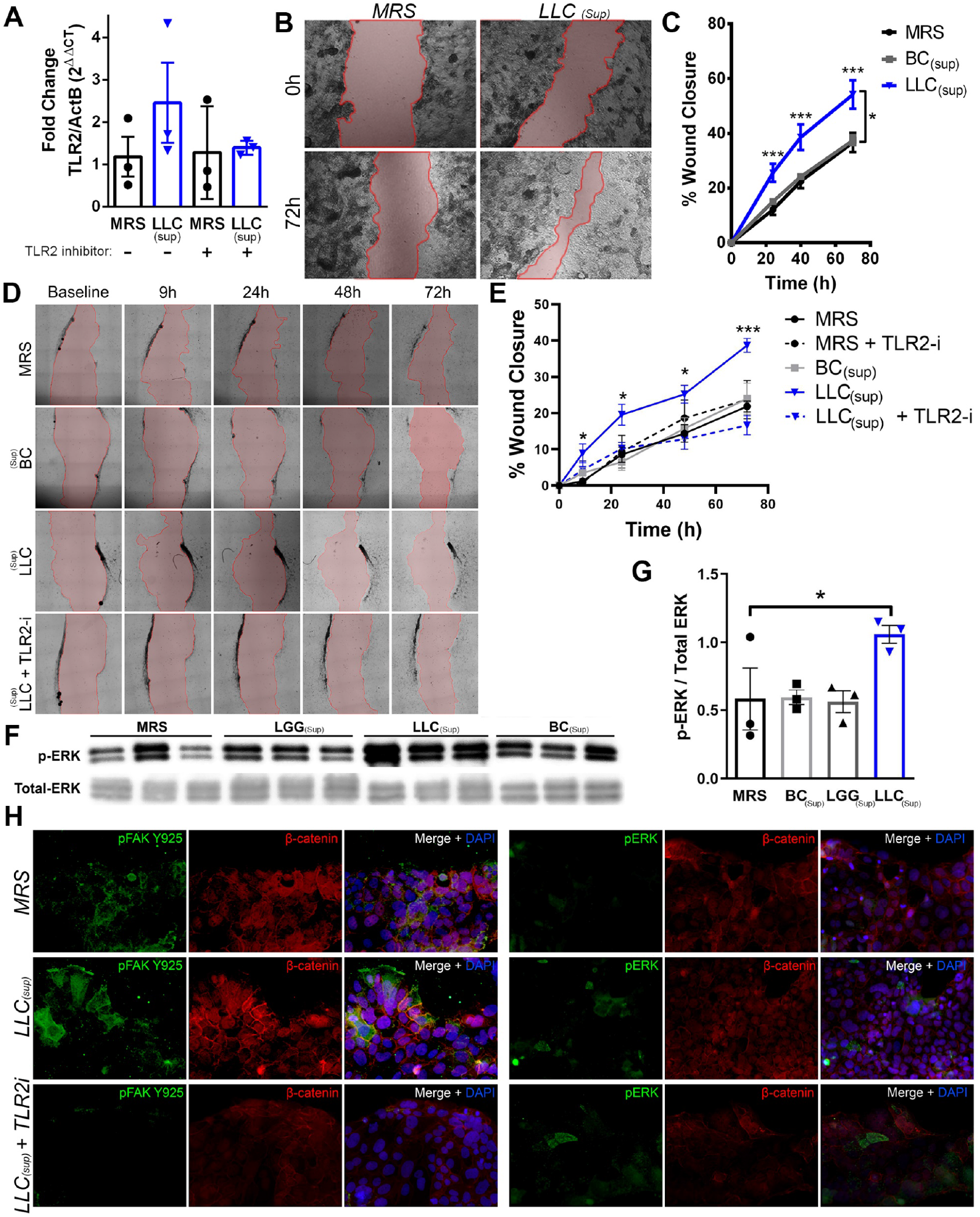
*L. lactis* subsp. cremoris promotes wound healing in cultured cells vitro through toll-like receptor 2 signaling. **(A**) RT-PCR of human colonic epithelium (T84 cells) after media supplementation with sterile filtered 1:100 LLC supernatant compared to sterile MRS bacterial culture media as a control, with and without TLR2 inhibitor. **(B)** Representative images of three independent scratch wound experiments of T84 monolayers treated with media supplemented 1:20 with either sterile MRS or LLC supernatant at 0 and 72 hours. **(C)** Quantitative analysis of scratch assays in **(D)** Representative images of three independent experiments of scratch wounds in human colonic epithelium (Caco2 cells) at 0, 9, 24, 48 and 72 hours of Caco2 monolayers treated with media supplemented with 1:20 LLC supernatants and TLR2 inhibitor daily **(E)** Quantitative analysis of scratch assays **(F)** Western Blot for p-ERK over total-ERK from T84 monolayers treated with MRS or LLC supernatant with or without TLR2 inhibitors **(G)** Densitometry of Western blot quantification for each group. Additionally, TLR2 inhibitors were utilized in a same approach as (D) and stained for p-FAK and p-ERK. **(H)** Representative images of Caco2 monolayers were seeded onto a glass coverslip, treated with LLC or MRS supernatants, with or without TLR2 inhibitors, subject to scratch wounds, fixed at 48h and stained by immunofluorescence for p-FAK and p-ERK. Data are represented as mean ± SEM, One way-ANOVA tests were used with multiple comparisons to determine the difference between groups *p < 0.05, **p < 0.01, ***p<0.001.***=p<.001. Scratch wound experiments performed N=3 in a 24 well plate in triplicate with each data point being a separate experiment (N=9 wells per group). N=3 for immunofluorescence and Western blot experiments.

### L. lactis subsp. cremoris supplementation induces TLR2 dependent activation of ERK and FAK to exert pro-restitutive effects on the colon

To investigate if TLR2 is required for *L. lactis* subsp. cremoris to elicit its beneficial influences on host health and disease, we supplemented the diet of C57BL/6 WT or TLR2-/- mice daily for 2 weeks with 1×10^8^ CFU *L. lactis* subsp. cremoris or vehicle control by oral gavage before repeated rounds of 2% DSS included in the drinking water of experimental groups for 28 days (**Figure 6A**). *L. lactis* subsp. cremoris supplementation mice significantly lowered disease activity in the colonic tissues of mice in a TLR2-dependent manner over the entire study period (**Figure 6B-E**). These data show that *L. lactis* subsp. cremoris dietary supplementation protects against gut injury induced by chronic Dextran Sodium Sulfate (DSS) in a TLR2-dependent manner. To confirm our observation of accelerated recovery following DSS-induced injury wound healing in DSS injured mice, we inflicted repeated biopsies on the colon of individual mice and quantitated healing rates by colonoscopy. 6-week-old C57BL/6 WT or TLR2-/- mice were supplemented with either 1×10^8^ CFU total *L. lactis* subsp. cremoris, *Bacillus cereus* (BC) or vehicle control for 7 days. Mice were then inflicted with 4 wounds at distances of 1, 2, 3 and 4cm into the colon using biopsy forceps (**Figure 6F)**. Measurement of the rate of healing of biopsy was assessed by colonoscopy at 48 hours and 96 hours after initial wounding revealed that mice supplemented with *L. lactis* subsp. cremoris had significantly accelerated healing rates in a TLR2-depenendet manner (**Figure 6G-H)**. To confirm our findings of *L. lactis* subsp. cremoris-induced activation of ERK and FAK signalling in cultured cells, we also assessed pathway signalling within biopsy wounds. Twelve-week-old WT C57BL/6 or TLR2-/- mice were given four wounds at distances of 1, 2, 3 and 4cm into the colon using biopsy forceps, then subjected to rectal gavage with 1×10^9^ CFU of *L. lactis* subsp. cremoris, or media control. After 30 mins, mice were sacrificed, and wound beds mounted in OCT for immunostaining. We detected markedly elevated levels of phospho-FAK within the wound beds of WT C57BL/6 supplemented with *L. lactis* subsp. cremoris in a TLR2-dependent manner (**Figure 6I)**. These data show that *L. lactis* subsp. cremoris supplementation promoted the activation of pathways that function in healing of biopsy wounds including the activation of FAK phosphorylation within biopsy wounds in a TLR2-dependent manner.

**Figure 6.**
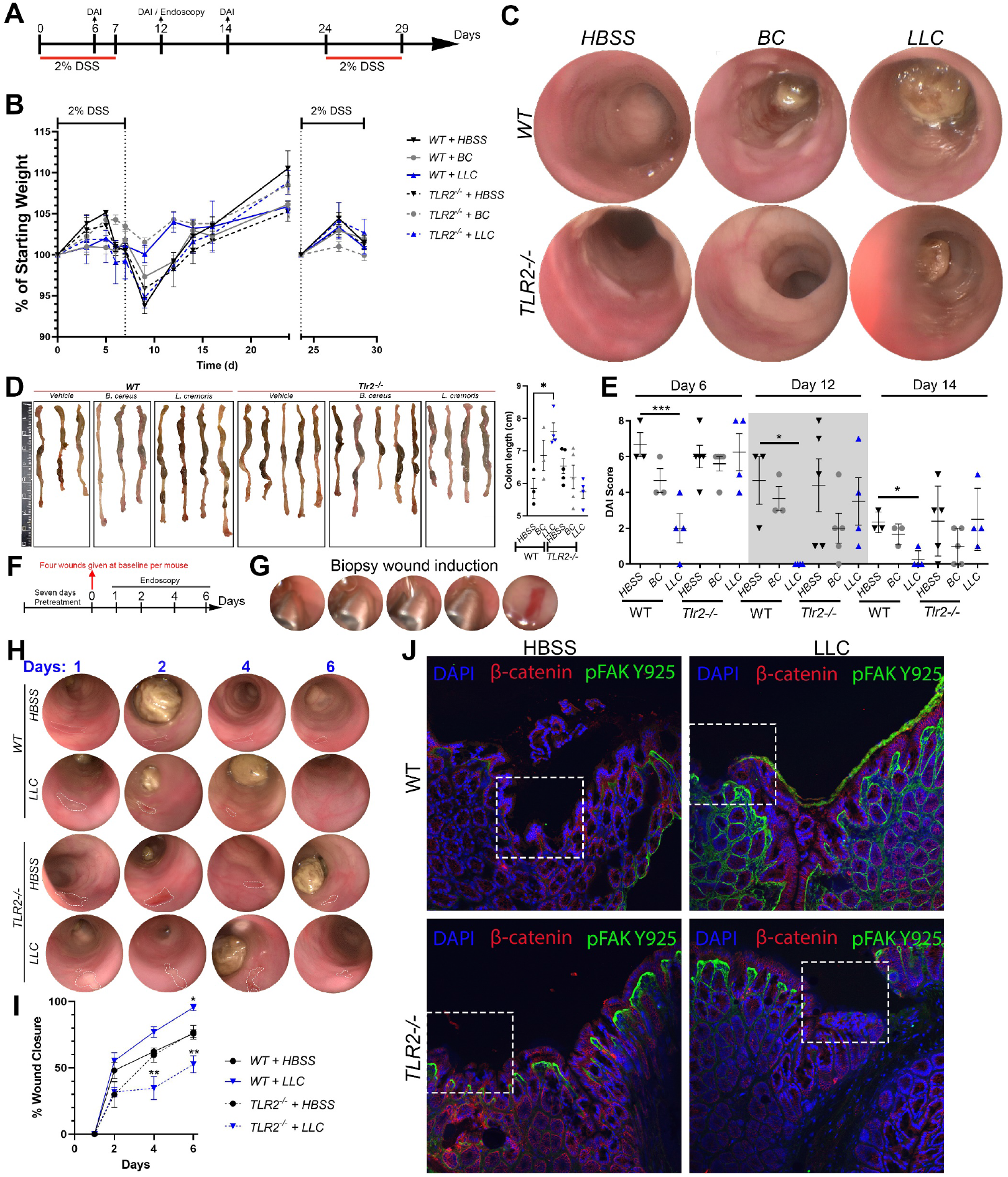
*L. lactis* subsp. cremoris induces TLR2 dependent activation of ERK and FAK to exert pro-restitutive effects on the colon. **(A)** Six-week-old male *Tlr2-/-* and WT C57Bl/6 mice were administered either 1 × 10^9^ CFU total Lactococcus lactis subsp. cremoris ATCC 19257 (LLC), Bacillus cereus (BC) or saline vehicle (HBSS) for 7 days pretreatment. 2% DSS was included in the drinking water (day 0) for 5 days in the indicated groups. Once all groups of mice had recovered to 95% of their original starting weights, 2% DSS was re-introduced into the drinking water for another 3 days. Weight was tracked in mice and probiotic or vehicle treatments were administered daily throughout. N=4-5 per group. **(B)** Mouse weight over time. **(C)** Representative colonoscopic images of median DAI mice from each group at day 12 **(C)** Representative images of colon length of mice and quantification of colon length of mice. N= 4-5. **(E)** Disease activity index (DAI) of mice at day 6, 12 and 14. **(F)** Twelve-week-old male TLR2-/- and WT C57BL/6 mice were administered either 1 × 109 CFU total Lactococcus lactis subsp. cremoris ATCC 19257 (LLC), Bacillus cereus (BC) or HBSS (vehicle control) for 7 days. Mice were given four wounds at distances of 1, 2, 3 and 4cm into the colon using biopsy forceps, then endoscopically examined at these distances into the colon every two days to measure wound closure rates. **(G)** Representative induction of a shallow wound into the mouse colon via endoscopic biopsy **(H)** Representative images from wounds n=3 wounds per mouse and n=3 mice per group. Wounds edges are indicated by dotted lines. **(I)** Wound area over time. **(J)** Twelve-week-old male TLR2-/- and WT C57Bl/6 mice were given four wounds at distances of 1, 2, 3 and 4cm into the colon using biopsy forceps, then subject to rectal gavage with 1×109 CFU of LLC or BC, or HBSS and sacrificed after 30 mins. Wound beds were mounted in OCT and immunostained for phospho-FAK or phospho-ERK. Representative images are shown, n=3 per group. Data are represented as mean ± SEM, One way-ANOVA tests were used with multiple comparisons to determine the difference between groups *p < 0.05, **p < 0.01, ***p<0.001.***=p<.001.

### *L. lactis* subsp. *cremoris* supernatants elicits improved survival in vitro and promotes proliferation and increased stemness in *ex vivo* enteroid cultures

TLR2 functions in the sensing of bacterial cell-wall components, such as lipoproteins from Gram-negative and Gram-positive bacteria, and lipoteichoic acid from Gram-positive bacteria. We showed that *L. lactis* subsp. cremoris does not protect Myd88-/- or TLR2-/- mice from gut injury. To further test the extent to which a factor released by *L. lactis* subsp. cremoris can elicit beneficial effects on cells, we collected filter sterilized supernatant from centrifuged cultures of this bacterium and included it in the growth media of cultured human T84 colonic epithelial cells. Measurements of lactate dehydrogenase (LDH) levels, which is is rapidly released into the cell culture supernatant when the plasma membrane is damaged as cells undergo apoptosis, necrosis, or other forms of cellular damage revealed significantly lower levels of LDH released by cultured T84 cells overlaid with *L. lactis* subsp. cremoris culture supernatant compared to controls. **(Figure 7A)**. These data are evidence that *L. lactis* subsp. cremoris does not induce any significant toxicity on cultured cells. In addition, we assessed the capacity of *L. lactis* subsp. cremoris filtered culture supernatant to promote tissue development. We generated ex vivo enteroids from C57BL/6 WT and *TLR2-/-* mice. At day 1 of growth, enteroids media was supplemented with filter sterilized cultured supernatants of *L. lactis* subsp. cremoris, *B. cereus*, or MRS bacterial culture media. Analysis of development by quantification of enteroid budding showed that enteroids overlayed with *L. lactis* subsp. cremoris supernatant had significantly increased growth (budding) after 6 days of enteroid growth compared to MRS control in a TLR2- dependent manner **(Figure 7B-D)**. Of note, *TLR2-/-* enteroid budding was significantly delayed in each experimental group irrespective of treatment. Finally, consistent with our results in cultured cells and in biopsy wounds, we also show that *L. lactis* subsp. cremoris filter sterilized culture supernatant markedly activated phospho-FAK and phospho-ERK in situ in enteroids after 6 days of growth **(Figure 7E)**. Together, these data show that *L. lactis* subsp. cremoris supernatants elicits no toxic influences in T84 cells and promotes proliferation and increased stemness in *ex vivo* enteroid cultures in a TLR2-dependent manner.

**Figure 7.**
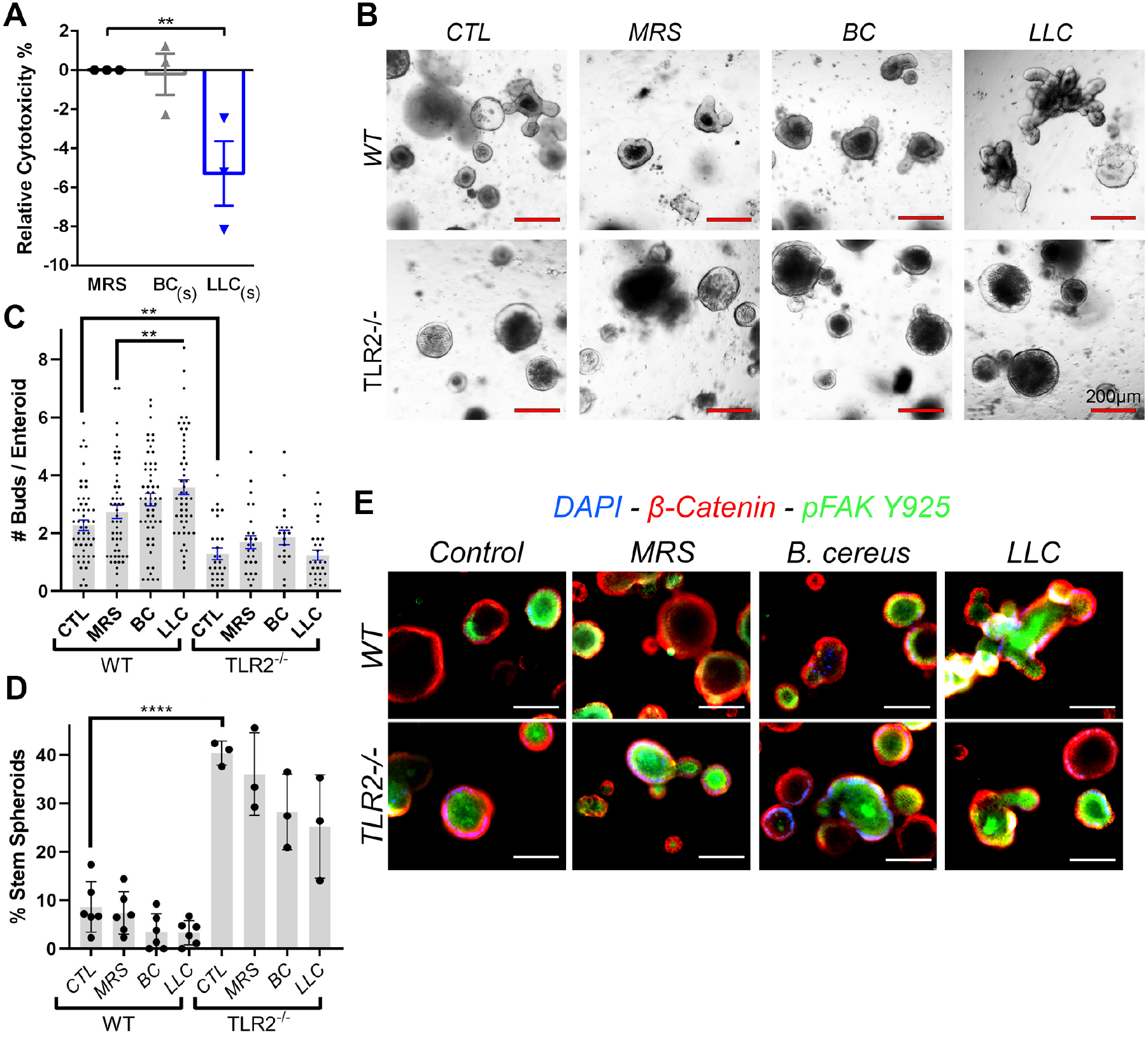
*L. lactis* subsp. cremoris supernatants elicits improved survival in vitro and promotes proliferation and increased stemness in *ex vivo* enteroid cultures. **(A**) LDH release assay for cytotoxicity of human colonic epithelium (T84 cells) after treatment with 1:100 LLC supernatants compared to non-probiotic bacteria Bacillus cereus (BC) supernatants and MRS media demonstrates that LLC confers improved survival in colonic epithelia. **(B)** Colonic crypts were isolated from WT or *TLR2-/-* mice and embedded in Matrigel. Representative bright field images of enteroids from WT and *TLR2-/-* mice at 6 days of growth, treated with MRS media (control) and BC or LLC supernatants at 1:100. **(C)** Buds per enteroid. Each data point is an average of buds counted in five enteroids (WT N=236-281 were counted per group, *TLR2-/-* N=108-131 were counted per group) **(D)** Ratio of stem spheroid proportion to total count after 6 days of growth post-plating, each data point represents the average per well, with N=43-187 colonoids counted per well **(E)** Immunofluorescent staining of phospho-FAK and phospho-ERK in situ in enteroids after 6 days of growth. Data are represented as mean ± SEM, One way-ANOVA tests were used with multiple comparisons to determine the difference between groups *p < 0.05, **p < 0.01, ***p<0.001.***=p<.001.

## Discussion

An ever-expanding body of evidence validates the concept that beneficial bacteria within the intestine positively influences health by stimulating gut homeostasis and promoting tissue restitution following injury. However, few of the molecular mechanisms whereby beneficial bacteria signal to sub-epithelial compartments to elicit their influence on the host have been recognized. In this study, we identified a set of genes induced by *L. lactis* subsp. cremoris in the colon that are associated with host tissue homeostasis and response to injury. We showed that *L. lactis* subsp. cremoris also elicited cytoprotection from colitis injury in mice in a Myd88 and TLR2-dependent mechanism. We showed that the culture supernatant from *L. lactis* subsp. cremoris was sufficient to induce accelerated wound healing and activation of ErK and FAK siganling in a TLR2-dependent manner. These data define a mechanimsm whereby *L. lactis* subsp. cremoris signals via the TLR2/MyD88-axis to confer beneficial infleucnes on the the gut epithelium.

These data contribute to the gradually expanding literature describing the effects of beneficial bacteria on host tissue. The focus of previous investigations from our group was the most widely studied strain of probiotic bacteria, namely *Lactobacillus rhamnosus* GG. We previously reported that *L. rhamnosus* GG facilitates intestinal stem cell turnover [36], wound restitution in the gut [43], mediates cytoprotective effects in the gut [38], as well as cell migration [42, 44]. We also showed that *L. rhamnosus* GG can prevent sex steroid depletion-induced bone loss resulting in decreased gut permeability and intestinal and systemic inflammation [45], as well as showing the role *L. rhamnosus* GG plays in stimulating bone formation by expanding the pool of Treg cells in the gut and the bone marrow [46]. Other reported mechanism of action of LGG are via the generation of the P40 and P75 proteins that belong to a family of secreted cell wall proteins that contain a carboxy(C)-terminal CHAP or NlpC/P60 superfamily domains. P40 was shown to prevent cytokine-induced epithelial damage and apoptosis [47–51]. Another very well-studied beneficial bacterium is *Bacteroides fragilis*, which released a bacterial polysaccharide (PSA) directs the cellular and physical maturation of the developing immune system [52]. *B. fragilis* was shown to be equally effective in preventing colitis and allergic encephalomyelitis (EAE) in murine models[53–55]. The immunomodulatory activities of PSA was shown to induce regulatory T cells to secrete IL-10, a potent anti-inflammatory cytokine that restrains pathogenic inflammation in the gut, as well as systemically in the brain [54, 56–58].

The limitations of our previous report on *L. lactis* subsp. *cremoris* was that the molecular mechanism whereby *L. lactis* subsp. *cremoris* confers its beneficial effects remained unknown [39]. We reported on the mechanisms whereby *Lactobacillus rhamnosus* GG elicits its beneficial response via the Nox1-dependent cellular generation of reactive oxygen species (ROS), which acts as a signaling molecule to transduce bacterial-induced signals to sub epithelial compartments [36, 38, 42, 43, 59–61]. Furthermore, *L. rhamnosus* GG is reported to be sensed by host epithelial cells via sensing by formyl peptide receptors (FPRs), which are G protein-coupled receptors and were first characterized in the transduction of chemotactic signals in phagocytes, and the mediation of host-defense and inflammatory responses by superoxide production [42, 43]. Because contact of *L. lactis* subsp. *cremoris* with epithelial cells does not activate detectable cellular ROS generation [39], we speculated that *L. lactis* subsp. *cremoris* elicited its beneficial effects by a different signaling mechanism to *L. rhamnosus* GG. The employment of methodologies such as RNAseq which measures transcript enrichment in tissues was instrumental in elucidating candidate functional elements that transduce the beneficial effects of *L. lactis* subsp. *cremoris* to the colonic tissue. The function of these elements, namely TLR2 and Myd88 were are confirmed using mice with knockout mutations that disrupt their function. These data highlight a contrast between the mechanism whereby two lactic acid bacteria, namely *L. lactis* subsp. *cremoris* and *L. rhamnosus* GG are sensed by host cells to elicit their beneficial effects.

Existing treatments for gut inflammation include oral dosing of corticosteroids, antibiotics, and immunomodulators by subcutaneous or intravenous administration [62]. The concept of employing microbial-based platforms as treatment to treat intestinal injury is an progressively accepted as a feasible approach [63]. Most existing beneficial bacteria are lactic acid bacteria from the lactobacilli genus [64–67]. Importantly, the characterization of *L. lactis* subsp. cremoris as an effective bacterium for limiting gut injury, identifies a beneficial lactococcus species with necessary features for therapy. *L. lactis* strains have been widely used in the production of dairy products, and consequently have a generally recognized as safe (GRAS) status. These desirable properties positively contribute to the prospect of using *L. lactis* subsp. *cremoris* as a therapeutic to treat intestinal ailments since potential toxological effects may be unlikely. Together, the innovative approaches of identifying a new and substantially more efficacious beneficial bacteria will yielded a considerable leap in improving the concept behind, and the usage of beneficial bacteria. Ultimately, these discoveries will exert a sustained and powerful influence by directly informing decisions related to data for product formulation, doses and timing for an Early Clinical Trial to substantiate the use of *L. lactis* ATCC 19257 as a therapeutic modality to accelerate recovery from, preserve remission, or prevent IBD.

## Materials and methods

### Experimental Mice

All experiments were done using C57BL/6 mice, TLR2 full body knockout mice (Stock No: 004650) and Myd88 knockout mice (Stock No: 009088) purchased from The Jackson Laboratories. Animal procedures were approved by the Institutional Animal Care and Use Committee of Emory University.

### Bacterial strains and culture preparation and administration of Bacteria

The following bacteria were purchased from the American Type Culture Collection (ATCC) (Manasas, VA): *Lactobacillus rhamnosus* GG ATCC 53103 and *Lactococcus lactis* subspecies cremoris ATCC 19257. All media was propagated according to instructions provided by the ATCC. *Bacillus cereus* was isolated by our research group [36]. Mice were gavaged with commensals as described previously [39]. Briefly *Lactococcus lactis* subspecies cremoris ATCC 19257, *Bacillus cereus*, or vehicle control, HBSS, were orally gavaged to mice every day in the mid-afternoon at a dose of 2×10^8^ CFUs at a volume of 100uL with control mice receiving 100uL of HBSS.

### Real-time PCR

Mouse colonic and intestinal tissue was frozen in liquid nitrogen and stored at −80°C until use. Tissues were homogenized for RNA extraction using TRIzol reagent (Invitrogen, Carlsbad, California, USA), and cleaned using RNeasy kit (Qiagen) total RNA clean up protocol. cDNA of these samples was synthesized from 1μg of total RNA using iScript Reverse Transcription Kit, and relative transcript levels were measured for each sample in triplicate by quantitative qRT-PCR using SYBR Green PCR Kit (Biorad), on a MyiQ™ Real time PCR system (Biorad). The data generated by qPCR assays were normalized using the average value of the HBSS treatment control group.

### Scratch Wound Assay

Wound healing assays were conducted on T84 cells using a modified protocol as previously described[68]. Briefly, T84 cells were grown as confluent monolayers in 12-well plates and scratched using a P200 tip. Cellular debris from the wound was removed by washing twice in sterile PBS. Cell culture media containing the TLR2i, C29 was included in the culture medium at 10uM final concentration. Wounds were photographed hourly for 24 h using a Zeiss Axiovert 200 M microscope with a Zeiss Incubator XL-3 489 attachment, maintaining cells at 37 °C/5% CO_2_. Cellular migration was measured using AxioVision v4.9.1 software (Carl Zeiss, Oberkochen, Germany) by measuring the size of the wound at each time point. Three individual measurements were made on each wound and averaged. Three independent experiments with a minimum of three replicates per experiment were pooled. Wound closure was calculated as percentage wound closure from *t* = 0.

### DSS-colitis model in mice

Groups of C57BL/6 mice were purchased from Jackson laboratory and maintained the Emory University Department of Animal Resources. Pure cultures of beneficial bacteria, including, or L. lactis cremoris, or *B. cereus* (2×10^9^ cfu total) were administered by oral gavage daily for indicated time periods, typicall between 7 and 14 days. Dextran Sodium Sulfate (DSS) was included in the drinking water of some groups of mice to a final concentration of 3% for acute DSS injury, or 2% for the chronic injury model. Body weights, and disease activity index were monitored daily until mice had lost 80% body weights (seven days), whereupon DSS was removed from the drinking water. Murine body weights, and disease activity index (DAI) were monitored daily for a further three days before mice were sacrificed and histological sections of the colon were prepared for disease activity. All animal experiments were approved by the Emory University institutional ethical committee and performed according to the legal requirements.

### Endoscope

All procedures and post-anesthesia care were performed in accordance within IACUC guidelines. Each mouse was anesthetized with 3% isoflurane and maintained with 1.5% isoflurane in an oxygen/air mixture by using a gas anesthesia mask. Body temperature was maintained during the procedure at 37°C with a homeothermic blanket (Harvard Apparatus, Holliston, MA). For monitoring of tumor burden, a high-resolution mouse video endoscopic system was utilized (Karl Storz Endoskope, Tuttlingen, Germany). The system consists of a miniature endoscope (1.9 mm outer diameter), a xenon light source, a triple chip camera, and an air pump for regulated inflation of the mouse colon (Karl Storz, Tuttlingen, Germany). The endoscopic procedure was viewed on a color monitor and digitally recorded for post-procedure review.

### Enteroid cultures

After washing the mouse colons and intestines (approx. 15cm of Jejunum) 15 times with cold phosphate buffered saline (PBS) without Ca2+/Mg2+, the isolated intestinal tissue was cut into ∼3-to 5-mm pieces and incubated in Gentle Cell Dissociation Reagent (Stem Cell Technologies, Vancouver, Canada). After removal of the Gentle Cell Dissociation Reagent, and resuspension with PBS with 0.1% BSA, the villi and mucus were removed with a 70-μm cell strainer (BD Biosciences, NJ). After being centrifuged at 8°C at 290 x g for 5 minutes, the collected crypts were resuspended in cold DMEM / F12 with 15mM HEPES and counted. 150 enteroid or colonoid crypts were plated in each well in fresh Matrigel at a ratio of 1:1. Passage of enteroids or colonoids derived from single cells was performed every 7-8 days. Post-passaging, half of the colonoids were stimulated with 100ng/ml of PEGylated leptin (Peptides International, KY) added to media 3 times weekly. Light microscopy was performed on cultures 2 days post passaging, with representative images selected after counting 6 images at 100x magnification from each group. Organoid counts were performed 2, 3- and 7-days pre-passaging, and at 7 days post-passaging, with small spheres consisting of enclosed sphere structures, large spheres having a defined ‘lumen”, budding spheres having one bud from the main structure and enteroids or colonoids having multiple buds.

### Antibodies and reagents

Colon samples for immunofluorescence were swiss-rolled and fixed in 10% formalin and then processed and embedded in paraffin. 5-micron sections were cut and underwent a deparaffination process followed by a citrate antigen retrieval step before being permeabilized in 0.5% triton-X 100 in PBS and blocked with 5% normal goat serum. Primary antibodies include, from Cell Signaling Technology (Danvers, MA), anti-Phospho-p44/42 MAPK cat#4370, anti-phospho-FAK (Tyr397) cat#3283. Samples were incubated in primary antibody overnight at 4ºC. Samples were then washed three times and incubated for 1 hour in fluorochrome-conjugated secondary antibody. Samples were again washed and incubated with DAPI at a concentration of 1:10000 in PBS for 5 minutes. Samples were mounted using Prolong diamond antifade and imaged on an Olympus FV1000 confocal microscope at 20X. At least 3 sections and three different images were taken for each sample. For immunoblot analysis, immunoreactive species were detected using the aforementioned anti-Phospho-p44/42 MAPK cat#4370, anti-phospho-FAK (Tyr397) cat#3283, followed by anti-rabbit HRP or anti-mouse HRP followed by visualization with ECL chemiluminescence detection reagent (GE Healthcare Biosciences Piscataway NJ). Experiments were conducted in triplicate and the average density of the band quantified using ImageJ software.

### Statistical Analysis

One-way ANOVA, with Bonferroni’s multiple comparisons test, was used for analysis of western blot flow cytometry and NTA data. Wound healing assay and invasion assay data were analysed using a Kruskal-Wallis test with Dunn’s multiple comparisons. Two-way ANOVA with Bonferroni’s multiple comparisons was used for analysis of proliferation data. Differences were considered statistically significant for P-values < 0.05. All analyses were performed using GraphPad Prism v6.0.4 software (GraphPad, San Diego, CA, USA). Data are presented as the mean ± standard error of the mean (SEM).

## Author Contributions

RMJ, CRN and JAO conceived and designed the experiments. CRN and RNK performed all experiments on enteroids. CRN, JAO, LL, and MB performed experiments on the murine animal model. RMJ, CRN and JAO wrote the manuscript.

## Acknowledgments

CRN is supported by an American Heart Association fellowship 19POST34370006. JAO is supported by the NIH through F31CA247415. RMJ is supported, in part, by a startup fund from The Emory University School of Medicine.

